# Tonic GABA_A_ conductance favors temporal over rate coding in the rat hippocampus

**DOI:** 10.1101/738369

**Authors:** Yulia Dembitskaya, Yu-Wei Wu, Alexey Semyanov

## Abstract

Synaptic plasticity is triggered by different patterns of neuronal network activity. Network activity leads to an increase in ambient GABA concentration and tonic activation of GABA_A_ receptors. How tonic GABA_A_ conductance affects synaptic plasticity during temporal and rate-based coding is poorly understood. Here, we show that tonic GABA_A_ conductance differently affects long-term potentiation (LTP) induced by different stimulation patterns. The LTP based on a temporal spike - EPSP order (spike-timing-dependent [st] LTP) was not affected by exogenous GABA application. Backpropagating action potential, which enables Ca^2+^ entry through N-methyl-D-aspartate receptors (NMDARs) during stLTP induction, was only slightly reduced by the tonic conductance. In contrast, GABA application impeded LTP dependent on spiking rate (theta-burst-induced [tb] LTP) by reducing the EPSP bust response and, hence, NMDAR-mediated Ca^2+^ entry during tbLTP induction. Our results may explain the changes in different forms of memory under physiological and pathological conditions that affect tonic GABA_A_ conductance.

## Introduction

A tonic conductance mediated by extrasynaptic GABA_A_ receptors has received significant attention over the last two decades (1–5). Often termed as the tonic current or the tonic inhibition, it thought to be a mechanism that decreases the excitability of specific cell populations in the brain (6–10). The tonic GABA_A_ conductance is set by concentrations of ambient GABA, the expression of extrasynaptic GABA_A_ receptors, the presence of endogenous and exogenous modulators of these receptors (8, 11–14). Endogenous modulators include neurosteroids, which concentrations in the brain change in puberty, pregnancy and the ovarian cycle (15–17). Corresponding changes in the tonic GABA_A_ conductance can promote stress, anxiety and depression (15, 16, 18, 19). One of the socially relevant exogenous factors augmenting the tonic GABA_A_ conductance is ethanol (20, 21). Hence, acute ethanol intake impairs synaptic plasticity, learning, and memory (22). In contrast, chronic ethanol abuse down-regulates GABAergic system (23).

Although the tonic GABA_A_ conductance is often referred to as the tonic inhibition, it is involved in far more complex neuronal computation than it would be expected from simple inhibitory action. The tonic conductance modulates neuronal gain during synaptic excitation in small-size cells (e.g., cerebellar granule cells) and neuronal offset in larger cells (e.g., hippocampal pyramidal neurons) (5, 24, 25). The tonic GABA_A_ mediated decrease in the cell input resistance reduces both the membrane time constant and the membrane length constant (26, 27). These constants influence the shape of EPSPs, their integration and EPSP-spike coupling in hippocampal interneurons and pyramidal cells (28, 29). In addition, the tonic GABA_A_ conductance modulates synaptic plasticity and brain rhythms (29–31).

Moreover, the tonic activation of extrasynaptic GABA_A_ receptors is not always inhibitory. Immature neurons have depolarizing reversal potential for GABA (E_GABA_), which becomes hyperpolarizing in the adult brain (32). Nevertheless, several types of mature neurons retain depolarizing E_GABA_: e.g., striatal projection neurons (33), hippocampal granule cells (34), suprachiasmatic nucleus neurons (35), cortical pyramidal neurons (36), vasopressin-secreting hypothalamic neurons (37), interneurons of hippocampus and amidgdala (38, 39) Low level of the tonic GABA_A_ conductance excites hippocampal interneurons by small depolarization recruiting voltage-dependent membrane conductances (40). High level of the tonic GABA_A_ conductance inhibits these neurons by shunting. The tonic activation of presynaptic GABA_A_ receptors depolarizes hippocampal mossy fiber boutons and increases the synaptic release probability (41).

Thus, the tonic GABA_A_ conductance influences cell excitability and integration of synaptic inputs depending on the cell size, E_GABA_ and the magnitude of this conductance. However, it remained unclear if the tonic GABA_A_ conductance differently affects synaptic plasticity triggered by different neuronal network dynamics. To address this issue, we compared the effect of tonic GABA_A_ conductance on stLTP induced by a pairing of synaptic input activation and postsynaptic cell spiking (temporal coding) and tbLTP induced by theta-bursts of presynaptic cell firing (rate coding).

## Results

### Tonic GABA_A_ conductance has no effect on stLTP but suppresses tbLTP

We recorded field EPSPs (fEPSP) in CA1 *stratum* (*str.) radiatum* of rat hippocampal slices in response to extracellular stimulation of Schaffer collaterals (Fig. 1A). Two experimental protocols were used to trigger LTP in CA3 - CA1 synapses: stLTP protocol (paired synaptic and antidromic stimulation repeated 100 times within 10 min) and tbLTP protocol (4 synaptic stimulations at 100 Hz repeated 10 times with 200 ms interval). Each protocol induced LTP which was entirely blocked by 50 μM DL-2-amino-5-phosphonovaleric acid (DL-APV), an NMDAR antagonist (Fig. 1B-G). An increase in tonic GABA_A_ conductance was simulated by bath application of 30 μM GABA. Due to efficient uptake we assumed acting concentration of GABA in low micromolar range expected under physiological conditions *in vivo*. GABA increased the membrane conductance in CA1 pyramidal neurons to 122 ± 6 % of baseline (*n* = 5, *p* = 0.02, one-sample *t*-test; Fig. S1), but did not significantly affect stLTP (fEPSP slope: 121 ± 6 % of baseline in control, 30 min after stLTP induction, *n* = 8; 125 ± 9 % of baseline in the presence of GABA, *n* = 7, *p* = 0.753, two-sample *t*-test; Fig. 1B-D). However, it reduced tbLTP approximately by half (fEPSP slope: 161 ± 13 % of baseline in control, 30 min after tbLTP, *n* = 7; 130 ± 7 % of baseline in the presence of GABA, *n* = 8, *p* = 0.041, two-sample *t*-test; Fig. 1E-G). These results suggest that although both types of LTP are NMDAR-dependent, they are differently affected by the tonic GABA_A_ conductance.

**Fig. 1.**
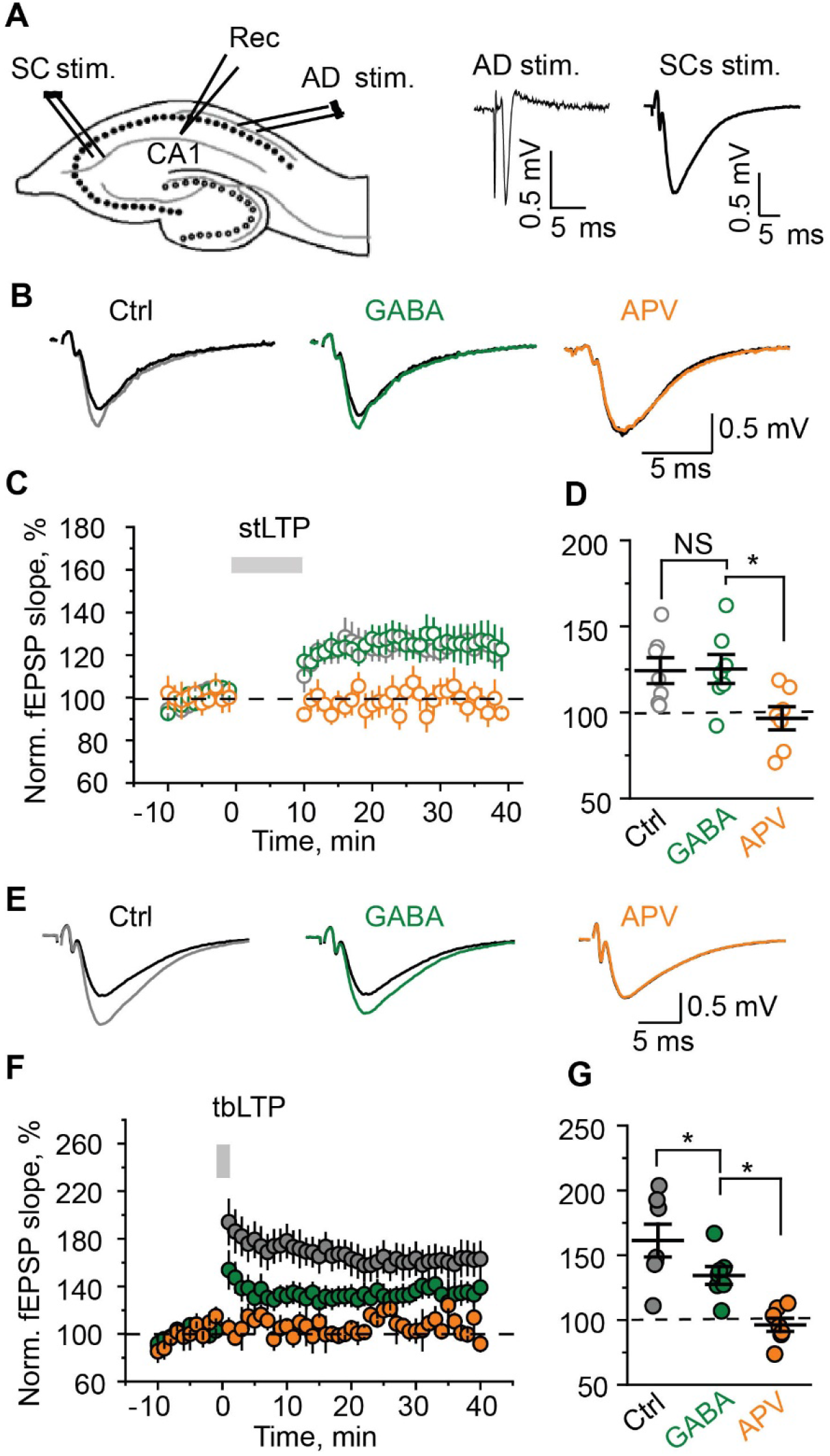
The tonic GABA_A_ conductance has no effect on stLTP but suppresses tbLTP. (A), *Left*, a schematic of hippocampal slice showing the location of stimulating and recording electrodes. The recording electrode was placed in CA1 *str.radiatum* (Rec), the first stimulating electrode on Shaffer collaterals (SC stim.), the second electrode in *str.oriens* for antidromic stimulation (AD stim.). *Right*, AD stim. induced antidromic population spike, while SC stim. induced fEPSP. stLTP was induced by the pairing of CS stim. and AD stim.; tbLTP was induced by bursts of SC stim. (B), Average fEPSPs 30-40 min after stLTP induction in control (grey), in the presence of GABA (green) and in the presence of APV (orange). Black traces are fEPSPs prior to stLTP induction. (C), The time-course of averaged and normalized fEPSP slope in control (grey), in the presence of GABA (green) and in the presence of APV (orange). Zero time point is the beginning of stLTP induction protocol (grey bar). (D), The summary data showing the magnitude of stLTP averaged during 30-40 min after induction; Control (Ctrl) - grey, GABA - green, APV - orange. (E), Average fEPSPs 30-40 min after tbLTP induction in control (grey), in the presence of GABA (green) and in the presence of APV (orange). Black traces are fEPSPs prior to tbLTP induction. (F), The time-course of averaged and normalized fEPSP slope in control (grey), in the presence of GABA (green) and in the presence of APV (orange). Zero time point is the beginning of tbLTP induction protocol (grey bar). (G), The summary data showing the magnitude of tbLTP averaged during 30-40 min after induction; Control (Ctrl) - grey, GABA - green, APV - orange. The data are presented as mean ± SEM. NS *p* > 0.05, \**p* < 0.05, two-sample *t*-test.

### Tonic GABA_A_ conductance does not affect AP properties

stLTP depends on voltage-dependent removal of the Mg^2+^-block of NMDARs by action potential (AP)-mediated depolarization. APs are regenerative events, and their amplitude depends on the activation of Na^+^ and K^+^ conductances, which are considerably larger than the tonic GABA_A_ conductance (42). Thus, the tonic GABA_A_ conductance should not be able to affect APs and, hence, NMDARs unblock during the stLTP-induction protocol. Nevertheless, previous reports have suggested that activation of extrasynaptic GABA_A_ receptors can influence AP waveform (43, 44). In contrast, we did not observe a significant change in the AP amplitude, the half-width or the threshold in the soma of CA1 pyramidal neurons in response to the GABA application (Table 1, Fig. 2).

**Table 1.**
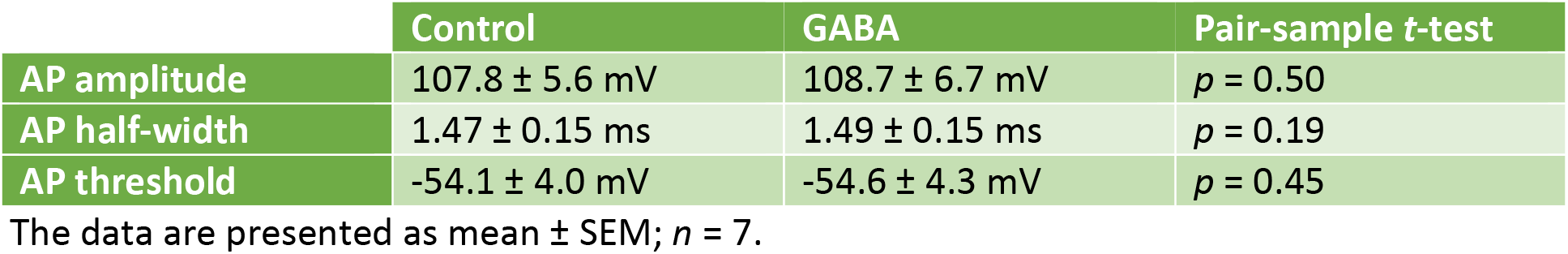
The tonic GABA_A_ conductance does not affect AP properties in the soma of CA1 pyramidal neuron.

**Fig. 2.**
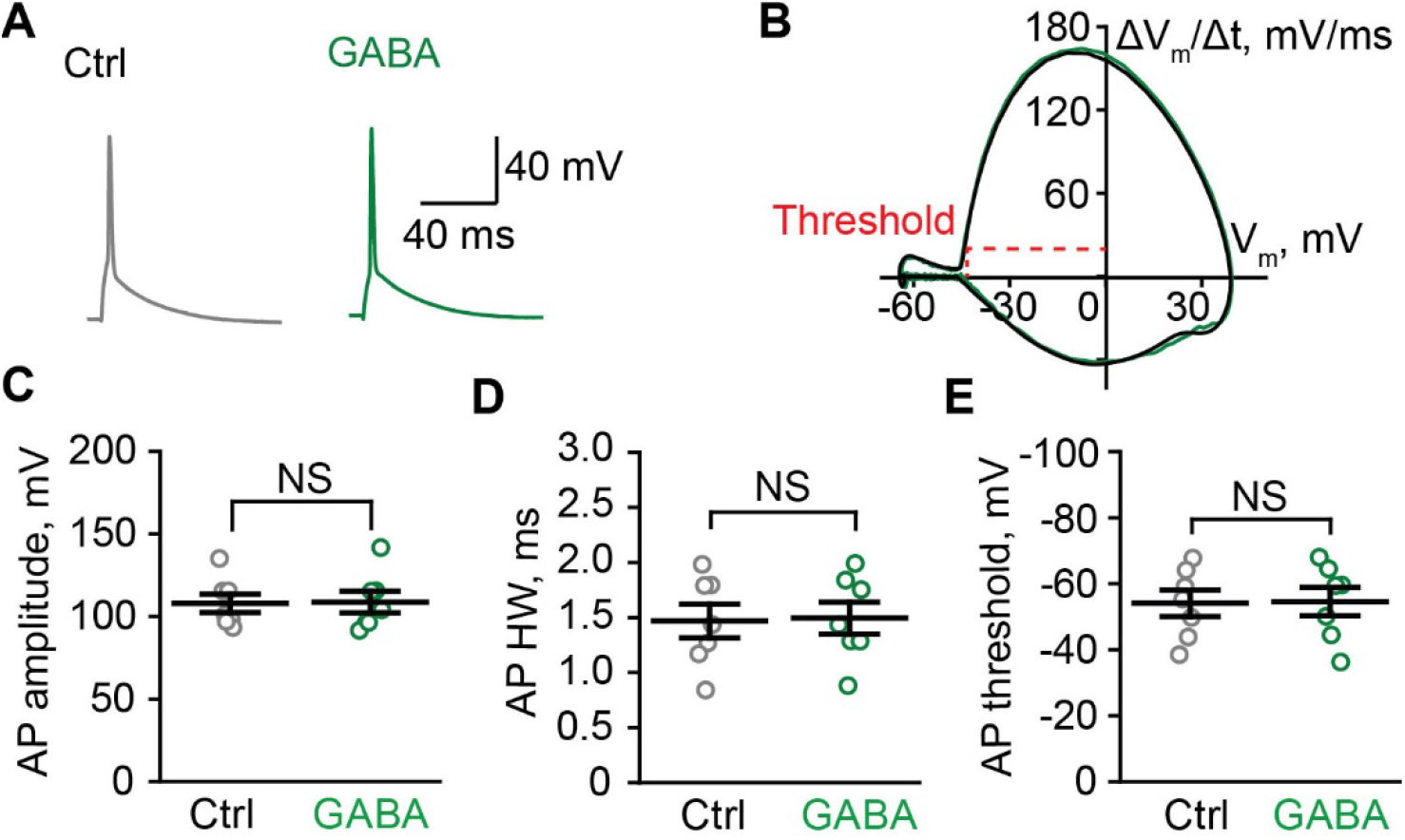
The tonic GABA_A_ conductance does not affect AP properties. (A), Sample APs in control (Ctrl, grey) and in the presence of GABA (green). (B), Phase plots of APs do not differ in control (grey) and in the presence of GABA (green). The threshold of AP was calculated as the membrane potential (V_m_) at AP velocity ΔV_m_/Δt = 20 mV/s. (C-E), The summary data showing no significant difference in the amplitude (C), the half-width (HW, D) and the threshold (E) of AP in control (grey) and in the presence of GABA (green). The data are presented as mean ± SEM. NS *p* > 0.05, pair-sample *t*-test.

### Tonic GABA_A_ conductance differently affects responses to bAP/EPSP pairing and EPSP burst in model pyramidal neuron

The properties of somatic APs are determined by the high density of Na^+^ and K^+^ channels in the nearby triggering zone. When the AP propagates along a dendrite, the density of voltage-dependent conductances decreases and such backpropagating AP (bAP) can be influenced by tonic GABA_A_ mediated shunting as a non-regenerative event (45, 46) but see (47). To address the effects of the tonic GABA_A_ conductance on bAPs in dendritic spines and thin dendrites we used a mathematical model suggested by Poirazi et al., 2003 (48). To induce a single EPSP activation of six synapses was simulated at proximal parts of apical oblique dendrites (Fig. 3A). A somatic current injection (1.7 nA, 2 ms) was simulated to trigger a bAP. The pairing of the EPSP and the bAP was used to mimic a single stimulation during the stLTP protocol. The somatic AP was triggered with 10 ms delay matching the bAP arrival with the peak of the EPSP. The bAP alone induced much smaller Ca^2+^ transient as compared to the EPSP (Fig. 3B). The pairing of the bAP with the EPSP induced a supralinear increase in the magnitude of the Ca^2+^ transient (49). Indeed, the tonic GABA_A_ conductance (1.7 mS/cm^2^) decreased the area under the curve (AUC) of the voltage response (ΔV_m_) induced by the bAP/EPSP pairing by approximately a quarter (Table 2; Fig. 3C). Hence, the amplitude of the Ca^2+^ transient induced by the bAP/EPSP pairing was reduced by a similar degree (Table 2; Fig. 3D,E). Subsequent blockade of NMDARs almost entirely suppressed the remaining Ca^2+^ transient (Table 2 GABA+APV; Fig. 3D,F). Notably, the blockade of NMDARs after introducing the tonic GABA_A_ conductance had no further effect on the AUC of ΔV_m_ induced by the bAP/EPSP pairing (Table 2; Fig. 3C). However, the blockade of NMDARs without introducing the tonic GABA_A_ conductance reduced the AUC of ΔV_m_ (Table 2; Fig. S2A). Such NMDARs blockade also suppressed the Ca^2+^ transient produced by the bAP/EPSP pairing (Table 2; Fig. S2B,C). Thus, most of Ca^2+^ entering the spine during the bAP/EPSP pairing is mediated by NMDARs. Approximately a quarter of this Ca^2+^ transient and of the voltage response is blocked by tonic GABA_A_ conductance.

**Table 2.** The tonic GABA_A_ conductance reduces the voltage response and the Ca^2+^ transient induced by the EPSP burst to a greater extent than these responses induced by the bAP/EPSP pairing.

|  | GABA | APV | GABA+APV |
| --- | --- | --- | --- |
| **bAP/EPSP pairing (mod)** |  |  |  |
| Dendrite $\Delta V_m$ AUC | 73.3 ± 0.4 % (6) | 87.5 ± 1.2 % (6) | 73.3 ± 0.7 % (6) |
| Peak $\Delta[Ca^{2+}]_i$ | 76.7 ± 1.8 % (6) | 6.8 ± 0.7 % (6) | 5.6 ± 0.5 % (6) |
| **EPSP burst (mod)** |  |  |  |
| Dendrite $\Delta V_m$ AUC | 48.8 ± 1.1 % (6) | 66.2 ± 0.2 % (6) | 44.6 ± 0.9 % (6) |
| Peak $\Delta[Ca^{2+}]_i$ | 53.8 ± 0.9 % (6) | 7.6 ± 0.6 % (6) | 0.0 ± 0.0 % (6) |
| **bAP/uEPSP pairing (exp)** |  |  |  |
| Soma $\Delta V_m$ AUC | 94 ± 2 % (7) | 94 ± 4 % (7) | 95 ± 3 % (7) |
| Peak $\Delta G/R$ | 86 ± 4 % (6) | 46 ± 4 % (6) | 43 ± 4 % (6) |
| **EPSP burst (exp)** |  |  |  |
| Soma $\Delta V_m$ AUC | 76 ± 6 % (6) | 76 ± 7 % (8) | 76 ± 5 % (6) |
| Peak $\Delta G/R$ | 71 ± 3 % (6) | 53 ± 5 % (6) | 41 ± 4 % (6) |

The responses were normalized to the control. GABA – the introduction of the tonic GABA<sub>A</sub> conductance, APV – blockade of NMDARs, GABA+APV – both. The data are presented as mean ± SEM (*n*). mod – model; exp – experimental data.

**Fig. 3.**
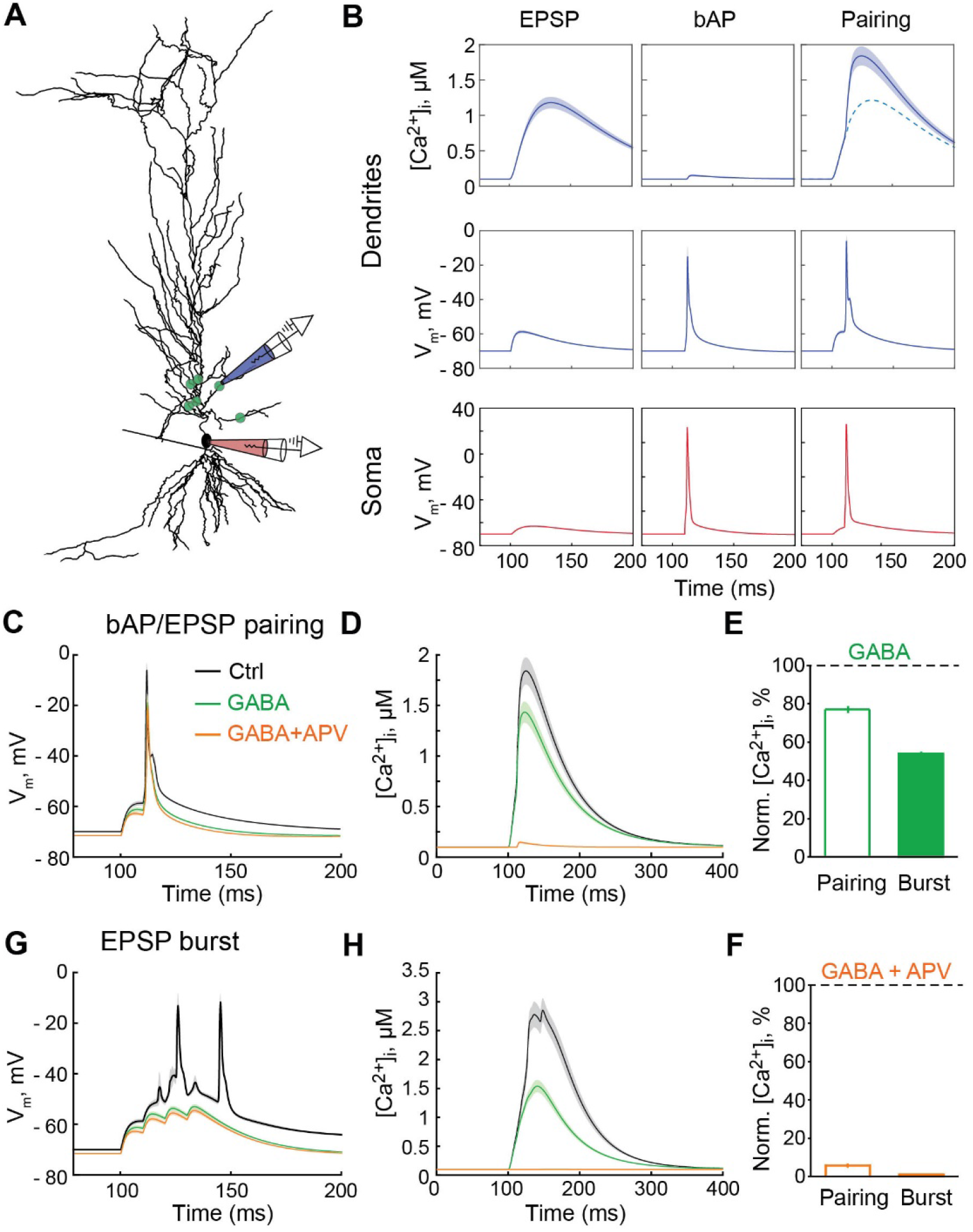
The tonic GABA_A_ conductance differently affects responses to the bAP/EPSP pairing and the EPSP burst in the model pyramidal neuron. (A), The synapse location (green dots) on the simulated pyramidal neuron. To electrodes indicate the places where the V_m_ was obtained: red – on soma, blue – on a second-order dendritic branc(H), The somatic electrode also indicates the place for current injection used to elicit bAP. (B), The Ca^2+^ transients in the dendrite (*top row*); V_m_ in the dendrite (*middle row*) and the soma (*bottom row*). *Left column* – the EPSP, *middle column* – bAP, *right column* – the bAP/EPSP pairing. [Ca^2+^]_i_ – intracellular Ca^2+^ concentration. (C), The voltage response to the bAP/EPSP pairing in control (Ctrl, grey), upon an increase in the tonic GABA_A_ conductance (GABA, green) and subsequent blockade of NMDARs (GABA+APV, orange). (D), The Ca^2+^ transient in response to the bAP/EPSP pairing in control (grey), upon an increase in the tonic GABA_A_ conductance (green) and subsequent blockade of NMDARs (orange). (E), The summary data are showing the effect of the tonic GABA_A_ conductance (GABA) on the amplitude of the Ca^2+^ transient induced by the bAP/EPSP pairing (empty bar) and the EPSP burst (filled bar). (F), The summary data are showing the effect of subsequent NMDAR blockade (GABA + APV) on the amplitude of the Ca^2+^ transient induced by the bAP/EPSP pairing (empty bar) and the EPSP burst (filled bar). (G), The voltage response to the EPSP burst in control (grey), upon an increase in the tonic GABA_A_ conductance (green) and subsequent blockade of NMDARs (orange). (H), The Ca^2+^ transient in response to the EPSP burst in control (grey), upon an increase in the tonic GABA_A_ conductance (green) and subsequent blockade of NMDARs (orange).

Next, we simulated the EPSP burst similar to that in the tbLTP protocol: 4 EPSP at 100 Hz were simulated at the same six synapses. The resulting EPSP burst triggered two somatic APs and a large Ca^2+^ transient in stimulated dendritic segments (Fig. 3G,H). The tonic GABA_A_ conductance reduced the EPSP burst below the threshold for AP generation and reduced the AUC of ΔV_m_ induced by EPSP burst approximately by half (Table 2; Fig. 3G). Consequently, the Ca^2+^ transient was also proportionally reduced (Table 2; Fig. 3F,H). Further blockade of NMDARs had a small effect on the EPSP burst voltage response, but completely abolished the Ca^2+^ transient (Table 2 GABA+APV; Fig. 3F-H). NMDAR blockade without introducing the tonic GABA_A_ conductance abolished one of two APs and, hence, reduced the voltage response (Table 2; Fig. S2D). It also largely suppressed the Ca^2+^ transient (Table 2; Fig. S2E,F). Thus, the model suggests that the tonic GABA_A_ conductance has a more profound effect on the voltage response and the Ca^2+^ transient induced by the EPSP burst than by the bAP/EPSP pairing.

### Tonic GABA_A_ conductance differently affects responses bAP/uEPSP pairing and uEPSP burst in CA1 pyramidal neuron

To confirm the model prediction, we performed two-photon Ca^2+^ imaging in dendritic spines (Fig. 4A). The dendritic Ca^2+^ transients (ΔG/R) induced by bAPs alone were not significantly affected by the GABA and subsequent 100 μM picrotoxin (GABA_A_ antagonist) application (GABA: 100.2 ± 8.2 % of control, *n* = 9, *p* = 0.98, one-sample *t*-test; picrotoxin: 96.0 ± 8.3 % of control, *n* = 9, one-sample *t*-test; Fig. S3). This finding suggests that the tonic GABA_A_ conductance did not affect bAP in imaged dendrites. This, however, does not rule out its action in more distal dendritic branches.

**Fig. 4.**
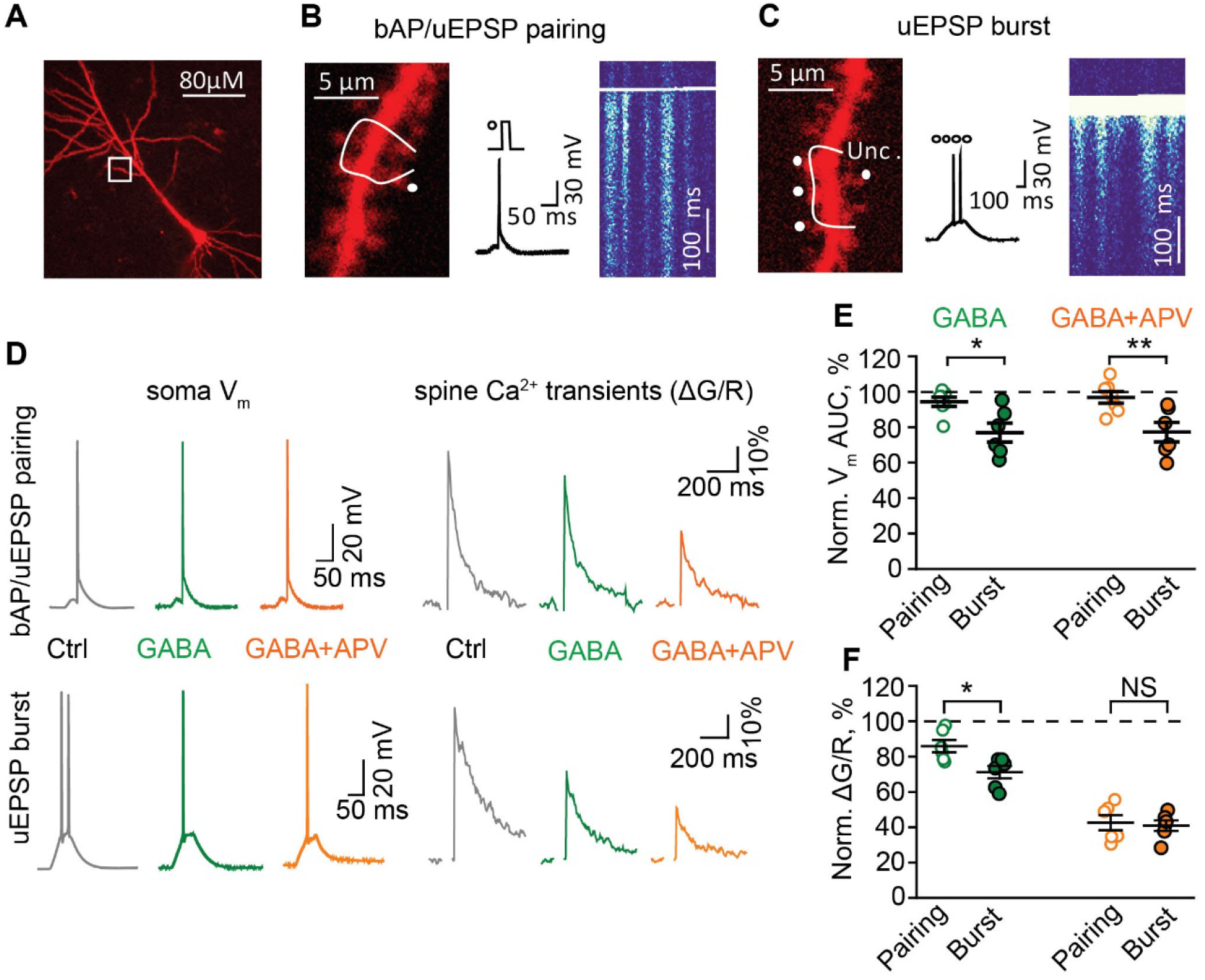
The tonic GABA_A_ conductance differently affects responses the bAP/uEPSP pairing and the uEPSP burst in CA1 pyramidal neuron. (A), CA1 pyramidal neuron filled with morphological tracer Alexa Fluor 594 (50 μM). The white box indicates a second-order dendritic branch used for Ca^2+^ imaging. (B), *Left* – zoomed in dendritic branch with a line scan trajectory (white line) and an uncaging spot (Unc., white dot); m*iddle* – the bAP/uEPSP pairing protocol (uncaging dot and current injection step) and corresponding V_m_ change in the soma; *right* – a line-scan image in Fluo-4F channel showing Ca^2+^ transients in dendritic spines and the shaft crossed by the scanning line, (C), *Left* – zoomed in dendritic branch with a line scan trajectory (white line) and four uncaging spots (Unc., white dots); m*iddle* – the uEPSP burst protocol (four uncaging episodes at 100 Hz) and corresponding V_m_ change in the soma; *right* – a line-scan image in Fluo-5F channel showing Ca^2+^ transients in dendritic spines and the shaft crossed by the scanning line, (D), The voltage responses in the soma (*left column*) and the Ca^2+^ transients in dendritic spines (ΔG/R, *right column*) triggered by the bAP/uEPSP pairing (*top row*) and the uEPSP burst (*bottom row*). *Grey traces* – control (Ctrl), *green traces* – GABA application, *orange traces* – subsequent APV application. (E), The summary data are showing the effect of GABA (green) and subsequent APV application (orange) on the area under the curve (AUC) of the voltage response to the bAP/uEPSP pairing (empty symbols) and the uEPSP burst (filled symbols). (F), The summary data are showing the effect of GABA (green) and subsequent APV application (orange) on the Ca^2+^ transients induced by the bAP/uEPSP pairing (empty symbols) and the uEPSP burst (filled symbols). The data are presented as mean ± SEM. NS *p* > 0.05, \**p* < 0.05, **p < 0.01, two-sample *t*-test.

The bAP/uncaging-induced (u)EPSP pairing was mimicked by single-photon spot photolysis of MNI-caged-L-glutamate (400 μM, glutamate uncaging) in front of a dendritic spine followed in 20 ms by an injection of depolarizing current through the patch pipette. The current was adjusted to trigger a single AP. 20 ms delay was introduced to match the bAP with the peak of the uEPSP (Fig. 4B). The uEPSP burst was induced by repeated glutamate uncaging (4 times at 100 Hz) at four synapses and was set to trigger two APs recorded in the soma in the control conditions (Fig. 4C). The GABA application did not have a significant effect on the somatic voltage response induced by the bAP/uEPSP pairing (*p* = 0.07, one-sample *t*-test; Table 2; Fig. 4D,E) because it dominated by somatic AP insensitive to the tonic GABA_A_ conductance. However, the Ca^2+^ transient induced by the bAP/uEPSP pairing was significantly reduced (*p* = 0.01, one-sample *t*-test; Table 2; Fig. 4D,F). In agreement with the model prediction, GABA reduced the number of APs induced by the uEPSP burst from 2 ± 0 to 1.1 ± 0.3 (*n* = 6, *p* = 0.031, pair-sample *t*-test, Fig. 4D). Hence, both the voltage response and the Ca^2+^ transient were significantly decreased (the AUC of ΔV_m_: *p* = 0.007, ΔG/R: *p* < 0.001, one-sample *t*-test; Table 2; Fig. 4D-F). Consistent with our hypothesis this decrease was significantly larger than the decrease of the responses induced by the bAP/uEPSP pairing (the AUC of ΔV_m_: *p* = 0.013, ΔG/R: *p* = 0.014, two-sample *t*-test; Table 2; Fig. 4D-F)

Subsequent blockade of NMDARs with DL-APV did not further decrease the voltage responses in both cases (Table 2 GABA+APV; Fig. 4D,E). However, the Ca^2+^ transients were reduced to a similar degree (*p* = 0.77, two-sample *t*-test; Table 2 GABA+APV; Fig. 4D,F). These results demonstrate that the Ca^2+^ transients induced both by the pAP/uEPSP pairing and by the uEPSP burst in dendritic spines are partially mediated by NMDARs and are sensitive to the tonic GABA_A_ conductance. However, it is not clear to what extent tonic GABA_A_ conductance affects the NMDAR-mediated and NMDAR-independent components of the Ca^2+^ transients. To address this issue, we reversed the order of the drug application: first, we applied DL-APV and then GABA. DL-APV had a small effect on the voltage response induced by the pAP/uEPSP pairing (*p* = 0.14, one-sample *t*-test) but significantly reduced the voltage response induced by the uEPSP burst (*p* = 0.008, one-sample *t*-test; *p* = 0.03, two-sample *t*-test for difference between stimulations; Table 2; Fig. S4). Nevertheless, NMDARs blockade reduced Ca^2+^ transients in both cases approximately by half (Table 2; *p* = 0.285, two-sample *t*-test; Fig. S4). Subsequent GABA application did not have a significant effect on the voltage responses, but further decreased the Ca^2+^ transients in both cases (to 39 ± 4 %, *n* = 6, for the pairing and 38 ± 3 %, *n* = 6, for the burst; *p* = 0.865, two-sample *t*-test; Fig. S4). This finding suggests that the tonic GABA_A_ conductance reduces both NMDAR-mediated and NMDAR-independent components of the Ca^2+^ transient. However, it affects differently only NMDAR-mediated Ca^2+^ transients.

## Discussion

We show that tonic GABA_A_ conductance suppresses tbLTP, but not stLTP. Both forms of plasticity require NMDARs, but these NMDARs are recruited by a different mechanism. At the baseline conditions, NMDARs are blocked by Mg^2+^ (50). Depolarization of the postsynaptic neuron removes this block and enables glutamate-bound NMDARs. While synaptic activity serves as a source of glutamate in both cases, the Mg^2+^ block is removed differently. In the case of tbLTP, the EPSP burst propagates to the soma and triggers APs, which propagate back and sum with the burst. The resulting depolarization unblocks NMDARs. The tonic GABA_A_ conductance suppresses the EPSP burst by shunting. Diminished EPSP burst is unable to trigger APs and recruits fewer NMDARs. Thus, tbLTP becomes smaller. In the case of stLTP, the voltage-dependent Mg^2+^ block of NMDARs is removed by bAP. The AP is a regenerative event mediated by significantly larger conductance than the tonic GABA_A_ conductance or a conductance underlying EPSP (42). Therefore, we did not detect any significant effect of the tonic GABA_A_ conductance on AP parameters (threshold, amplitude, HW) recorded in the soma. Nevertheless, this finding does not reflect the effect of the tonic GABA_A_ conductance on bAP in the dendrite. Dendrites have a lower density of Na^+^ and K^+^ channels than axon trigger zone which determines the shape of somatic AP (51–53). However, the bAP-induced Ca^2+^ transients recorded with two-photon imaging were not significantly affected by GABA application. As a result, the Ca^2+^ entry through NMDARs during the bAP/EPSP pairing was decreased by tonic GABA_A_ conductance significantly less than during the EPSP burst. Consequently, no significant decrease in stLTP magnitude was detected in contrast to tbLTP.

tbLTP can be linked to the cell firing during theta rhythms, which is important for the encoding and retrieval of space and time-related information (54–56). stLTP can occur when the neuron receives synaptic input from many presynaptic cells and need to choose which input is more relevant (57). Although a relevance of stLTP as a general model for synaptic plasticity is sometimes put into question (58, 59), this phenomenon has been demonstrated in different species and brain regions *in vitro* and *in vivo* (60–64). stLTP also has a broad appeal in computational neuroscience (65–69). Our finding that stLTP is more resistant to the tonic GABA_A_ conductance than tbLTP may provide further insights on how activity-dependent accumulation of ambient GABA can affect the brain computations, learning, and memory. It also can be considered in the context of physiological/pathological processes controlling the magnitude of the tonic GABA_A_ conductance: densities and properties of extrasynaptic GABA_A_ receptors; GABA release and clearance.

The tonic GABA_A_ conductance is mediated by multiple and plastic GABA_A_ receptors (13). The change in the magnitude of the tonic conductance due to modifications of receptor composition/density has been reported during development and pathological conditions (2). Indeed, both increased levels of the tonic conductance and attenuated LTP were reported in animal models of epilepsy (13, 70). In Alzheimer’s disease elevated concentrations of extracellular GABA occur due to GABA production and release by astrocytes (71). The increased tonic GABA_A_ conductance suppresses LTP in the hippocampal dentate gyrus in this disease.

Ambient GABA concentration builds up due to GABA spillover and, thus, reflects neuronal activity (11, 12). Neuronal activity can also be converted into an increase in extracellular GABA by astrocytes (72–74). Synaptically released glutamate is taken by astrocytic transporters along with Na^+^ with a 1:3 ratio. Intracellular Na^+^ accumulation reverses Na^+^-dependent GABA transporters which start to move GABA to the extracellular space.

High level of neuronal activity is also required for tbLTP induction (54). LTP induction in many synapses could potentially lead to excessive excitability of the brain, seizures, and excitotoxicity. However, an accompanying increase in extracellular GABA reduces both excitability and magnitude of tbLTP. Thus activity-dependent elevation of ambient GABA can serve as a protective mechanism to maintain the balanced level of brain excitation. This phenomenon is reminiscent of homeostatic downregulation of individual cells excitability by up-regulation of *h*-channels following LTP induction (75, 76). On the other hand, stLTP does not require a high rate of presynaptic firing but depends on the temporally correlated occurrence of synaptic input and postsynaptic AP. Thus, the induction of stLTP does not have to be associated with significant activity-dependent accumulation of extracellular GABA. In fact, tonic GABA_A_ conductance can be event beneficial for stLTP. stLTP is sensitive to spike jitter (77). Tonic GABA_A_ conductance can improve stLTP by reducing spike jitter through decreasing membrane time constant (28).

We conclude that brain states and activity increasing the tonic GABA_A_ conductance suppress rate coding (tbLTP) but not temporal coding (stLTP) in the hippocampus. This phenomenon may have important implications for overall brain computations, learning, and memory.

## Materials and Methods

### Hippocampal slice preparation

Transverse hippocampal slices were prepared from 3- to 5-week-old Sprague Dawley rats in accordance with RIKEN regulations. Animals were anesthetized with 2-bromo-2-chloro-1,1,1-trifluoroethane (halothane) and decapitated. The brain was exposed and cooled with ice-cold solution containing (in mM): 75 sucrose, 87 NaCl, 2.5 KCl, 0.5 CaCl_2_, 1.25 NaH_2_PO_4_, 7 MgCl_2_, 25 NaHCO_3_, 1 Na-ascorbate, and 11 D-glucose. Hippocampi from both hemispheres were isolated and placed in an agar block. Transverse slices (350-400 μm) were prepared with a vibrating microtome (Microm HM 650V, Thermo Fisher Scientific Inc., USA) and left to recover for 20 min at 34°C and then for 40 min in an interface chamber with storage solution containing (in mM): 127 NaCl, 2.5 KCl, 1.25 NaH_2_PO_4_, 2 MgCl_2_, 1 CaCl_2_, 25 NaHCO_3_, and 25 D-glucose. The slices were then transferred to the recording chamber and continuously perfused with recording solution containing (in mM): 127 NaCl, 2.5 KCl, 1.25 NaH_2_PO_4_, 1 MgCl_2_, 2 CaCl_2_, 25 NaHCO_3_, and 25 D-glucose at 34°C. All solutions were saturated with 95% O_2_ and 5% CO_2_. Osmolarity was adjusted to 295 ± 5 mOsm. 5 μM CGP52432 and 400 μM S-MCPG were routinely added to the solution to block GABAB and metabotropic glutamate receptors, respectively. Cells were visually identified under infrared DIC using an Olympus BX-61 microscope (Olympus, Japan).

### Electrophysiology

The glass electrode, with the resistance of 3 - 5 MΩ, filled with the extracellular solution was placed in *str.radiatum* for field potential recordings (Fig. 1A). Synaptic responses were evoked by stimulation with two bipolar stainless-steel electrodes (FHC, Bowdoinham, ME, USA). Theta-burst stimulation (TBS) of Schaffer collaterals was done with the electrode was placed in the *str. radiatum* at a distance more than 200 μm from the recording site to induce tbLTP. TBS stimulation consisted of 10 bursts with 200 ms inter-burst interval; each burst consisted of 4 pulses with the duration 0.2 ms and at frequency 100 Hz. stLTP was induced with one electrode placed in *str.radiatum* to trigger glutamate release from Schaffer collaterals (SC stimulation), and another electrode placed in *str.oriens* to trigger antidromic AP of postsynaptic CA1 pyramidal neurons (AD stimulation). Paired SC and AD stimulation were performed with a 10 ms delay of AD stimulation and repeated each 6 s for 10 min Stimulus strength was adjusted to 30-50% of the maximal amplitude of fEPSP. The slope of fEPSP was measured for data analysis.

Whole-cell recordings in CA1 pyramidal neurons were obtained using patch electrodes filled with a solution containing (in mM): 130 KCH_3_SO_3_, 8 NaCl, 10 HEPES, 10 Na_2_-phosphocreatine, 4 Na_2_ATP, 0.4 NaGTP, 3 L-ascorbic acid (pH adjusted to 7.2 with KOH, osmolarity to 290 mOsm) and with resistance of 3 - 5 MΩ. The recording solution also contained the morphological tracer Alexa Fluor 594 (50 μM, red channel) and Ca^2+^-sensitive dye Fluo-4F (250 μM, green channel) for stLTP, and Fluo-5 (300 μM, green channel) for tbLTP. The different dyes were selected because the baseline Ca^2+^ transients were significantly different for two types of stimulation and the dyes were selected to ensure that response was within the dynamic range.

APs were induced by somatic current injections (2 ms, 1-2 nA). The threshold of AP was calculated from phase portraits as the value of membrane potential (V_m_) when ΔV_m_/Δt was 20 mV/ms. Input resistance was calculated from the voltage response to 500 ms steps of current injections from −50 to +50 pA with a step of 10 pA (the slop of I-V curves). The series resistance was usually <20 MΩ, and data were discarded if it changed by more than 20 % during the recording. The series resistance was compensated with “bridge balance” function in current-clamp mode.

The signals were obtained with patch-clamp amplifier Multiclamp 700B (Molecular Devices, USA), filtered at 4 kHz and digitized at 10 kHz with NI PCI-6221 card (National Instruments, USA). The data were visualized and stored with software WinWCP (supplied free of charge to academic users by Dr. John Dempster, University of Strathclyde, UK).

### Two-photon imaging

Cells were filled with the dyes for at least 20 min before imaging to ensure dye equilibration. Two-photon Ca^2+^ imaging was performed with a two-scanner FV1000-MPE laser-scanning microscope (Olympus, Japan) equipped with a mode-locked (<140 fs pulse width) tunable 720–930 nm laser Chameleon XR (Coherent, USA). Both dyes were excited at 830 nm light wavelength, and their fluorescence was chromatically separated and detected with two independent photomultipliers (PMTs). We used bright Alexa Fluor 594 emission to identify oblique apical dendrites (about 150 mm from the soma) and their spines. Line-scan imaging was performed to record Ca^2+^ signals in the dendritic shaft and one to three spines. Imaging was synchronized with electrophysiological recordings. At the end of each recording, we tested that the Ca^2+^ transients were below the dye saturation level, which was achieved by prolonged somatic depolarization causing cell firing and Ca^2+^ buildup in the neurons. The changes in baseline Ca^2+^ level were monitored as the ratio between baseline Fluo-4/Fluo-5F and Alexa Fluor 594 fluorescence. If this ratio increased during the experiment by more than 20%, the recordings were discarded. The dark noise of the PMTs was collected when the laser shutter was closed in every recording.

### Glutamate uncaging

Bath application of 4-methoxy-7-nitroindolinyl-caged-L-glutamate (MNI-caged-L-glutamate; 400 μM) was used to ensure even distribution of the compound in the sample. Single-photon uncaging was carried out using 5–10 ms laser pulses (405 nm diode laser; FV5-LD405; Olympus) with “point scan” mode in Fluoview software (Olympus, Japan). The uncaging spots were usually positioned at the edge of spine heads of imaged dendrites. The Ca^2+^ transients were measured in response to (1) single bAP induced by somatic current injection; (2) “bAP/uEPSP pairing” when single bAP followed by uncaging at single dendritic spine with 20 ms delay (the strength of uncaging was adjusted to trigger 2-3 mV uEPSP); (3) “uEPSP burst” when uncaging was done simultaneously at 4 dendritic spines four times at 100 Hz (the strength of uncaging was adjusted to trigger two APs).

### Drugs and chemicals

All drugs kept frozen at −20°C in 100 to 200 μl 1000x concentration aliquots (stock solutions). 3-[[(3,4-Dichlorophenyl)methyl]amin-o]propyl] diethoxymethyl) phosphinic acid (CGP 52432), (S)-α-Methyl-4-carboxyphenylglycine (S-MCPG), γ-aminobutyric acid (GABA), D-(-)-2-Amino-5-phosphonopentanoic acid (APV), picrotoxin (PTX), 4-methoxy-7-nitroindolinyl-caged L-glutamate (MNI-glutamate) were purchased from Tocris Cookson (UK). Alexa Fluor 594, Fluo-4F, Fluo-5F were obtained from Invitrogen (USA).

### Data Analysis

Electrophysiological data were analyzed with WinWCP and Clampfit (Molecular Devices, USA). Imaging data were analyzed using FluoView (Olympus, Japan), ImageJ (a public domain Java image processing program by Wayne Rasband), and custom software written in LabView (National Instruments, USA). Statistical analysis was performed using Excel (Microsoft, USA) and Origin 8 (OriginLab, USA). The Ca^2+^ transient were represented as ΔG/R: (G_peak_ − G_baseline_)/(R_baseline_ − R_dark noise_); the baseline Ca^2+^ level was estimated as G/R, (G_baseline_ − G_dark noise_)/(R_baselin_ − R_dark noise_); where G is the Fluo-4F/Fluo-5F fluorescence, and R is Alexa Fluor 594 fluorescence. G _baseline_ and R _baseline_ are averaged fluorescence levels 50 – 100 ms before the stimulation. G_peak_ is mean fluorescence 30–40 ms after the stimulation. G_dark noise_ and R_dark noise_ are the dark currents of the corresponding PMTs. For illustration purposes, single traces were processed by five-point moving average, and then four to five sequential traces were averaged. The statistical significance was tested using a paired-sample or two-sample *t*-test when appropriate. The data are presented as mean ± SEM; n-number designates the number of recordings.

### Mathematical modeling

Simulations were performed with the NEURON 7.1 simulation environment (78). A biophysically detailed CA1 hippocampal pyramidal cell model was modified from the study by Poirazi et al., 2003 (48). The implemented membrane mechanisms included location-dependent R_m_ (membrane resistance) and R_a_ (axial resistance). The model included sodium, delay rectifier-, A-type, M-type, Ca^2+^-activated potassium, and *h*-type conductances; as well as L-, R-, and T-type voltage-dependent Ca^2+^ channels (48). The excitatory synaptic conductance was composed of AMPA and NMDA receptors (gAMPA and gNMDA): gAMPA was represented by a double exponential function with τ_rise_ of 1 ms, τ_decay_ of 12 ms (79); gNMDA was implemented from the study by Kampa et al., 2004 (80). Six excitatory synapses were placed at six different apical oblique dendrites (apical dendrite 5, 8, 10, 18, 113, 118) within the *str.radiatum*. gAMPA and gNMDA were set to generate a 10 pA somatic EPSC with a physiological NMDA/AMPA charge ratio in each excitatory synapse (79). The tonic GABAergic conductance (gGABA) with the outward-rectifying property was adapted from the study by Pavlov et al., 2009 (24) and was set to increase membrane conductance by 25% (1.7 mS/cm^2^) with E_GABA_ = −75 mV to mimic experimental condition in Fig. S1. The initial resting membrane potential of simulated neurons was set at −70 mV, and the simulation temperature was 34°C.

Membrane potential and intracellular Ca^2+^ concentration were simulated during the bAP/EPSP pairing and the EPSP burst. For the bAP/EPSP pairing activation of 6 excitatory synapses was followed by somatic current injection (1.7 nA, 2 ms) to trigger a single AP with 20 ms delay; for the EPSP burst, six excitatory synapses were activated four times at 100 Hz; To mimic in DL-APV and in PTX applications gNMDA and gGABA were set to zero, respectively.

## Acknowledgments

The authors are grateful to Dr Tanja Brenner for performing preliminary experiments on the project.

