## Supplementary information for "Tonic GABA_A_ conductance favors temporal over rate coding in the rat hippocampus"

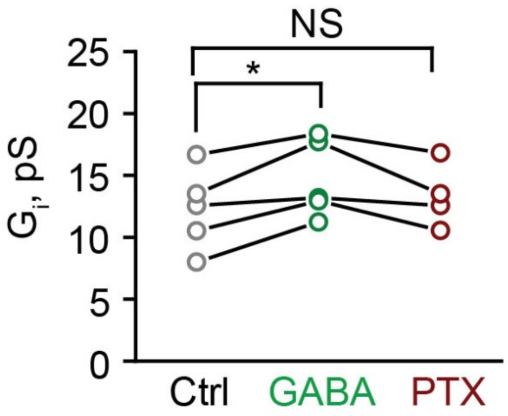


**Fig. S1.**

GABA increases cell input conductance. The input conductance ($G_i$) was increased by GABA application (green). The effect was reversed by $GABA_A$ receptor blocker picrotoxin (PTX, wine). Notably, PTX failed to significantly decrease the $G_i$ below the baseline level, which is consistent with the negligible baseline tonic $GABA_A$ conductance in CA1 pyramidal neurons in slices (1, 2).

The data are presented as mean ± SEM. *$p < 0.05$, NS $p > 0.05$, pair-sample *t*-test.

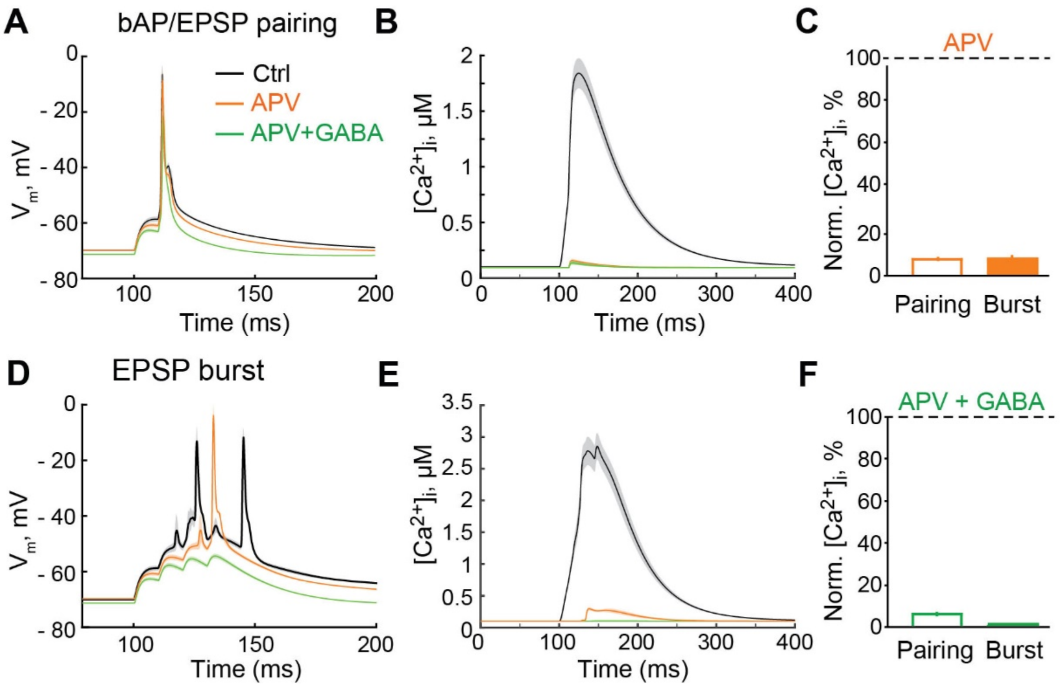


**Fig. S2.**

GABA effect on NMDAR-independent $Ca^{2+}$ transients in the model neuron. (A), Change in $V_m$ in response to the bAP/EPSP pairing in control (Ctrl, black), upon blockade of NMDARs (APV, orange) and subsequent increase of the tonic $GABA_A$ conductance (GABA + APV, green). (B), The $Ca^{2+}$ transient in response to the bAP/EPSP pairing in control (black), upon blockade of NMDARs (orange) and subsequent increase of the tonic $GABA_A$ conductance (green). (C), The summary data are showing the effect of NMDARs blockade (APV, orange) and subsequent increase in the tonic $GABA_A$ conductance (GABA + APV, green) on the amplitude of the $Ca^{2+}$ transient induced by the bAP/EPSP pairing. (D), The summary data are showing the effect of NMDARs blockade (APV, orange) and subsequent increase in the tonic $GABA_A$ conductance (GABA + APV, green) on the amplitude of the $Ca^{2+}$ transient induced by the EPSP burst. (E), Change in $V_m$ in response to the EPSP burst in control (Ctrl, black), upon blockade of NMDARs (APV, orange) and subsequent increase in the tonic $GABA_A$ conductance (GABA + APV, green). (F), The $Ca^{2+}$ transient in response to the EPSP burst in control (black), upon blockade of NMDARs (orange) and subsequent increase of the tonic $GABA_A$ conductance (green).

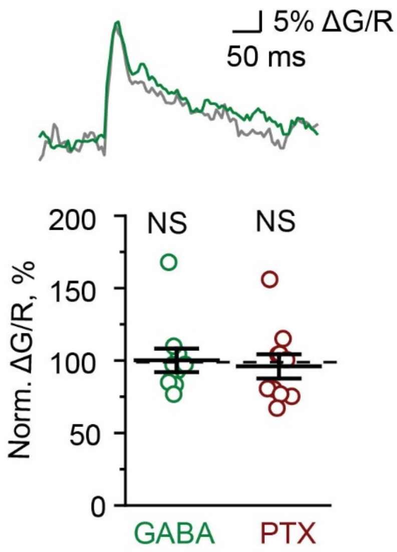


**Fig. S3.**

GABA does not significantly affect the bAP-induced $Ca^{2+}$ transients. The amplitude of the $Ca^{2+}$ transients (ΔG/R) in dendritic spines does not change after application of GABA (green) and subsequent application of 100 µM picrotoxin (PTX, wine). Sample traces: black – control; green – GABA.

The data are presented as mean ± SEM. NS $p > 0.05$, one-sample *t*-test.

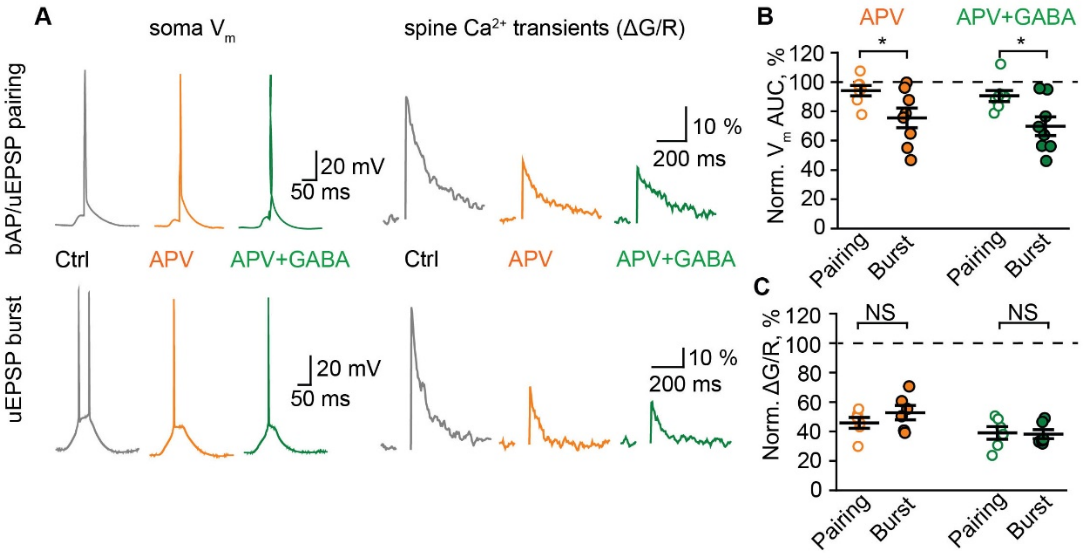


**Fig. S4.**

GABA does not significantly affect NMDAR-independent $Ca^{2+}$ transients. (A), The voltage responses in the soma (*left column*) and the $Ca^{2+}$ transients in dendritic spines (ΔG/R, *right column*) triggered by the bAP/uEPSP pairing (*top row*) and the uEPSP burst (*bottom row*). *Grey traces* – control (Ctrl), *orange traces* – APV application, *green traces* – subsequent GABA application. (B), The summary data are showing the effect of APV (orange) and subsequent GABA application (green) on the area-under-the-curve (AUC) of voltage response to the bAP/uEPSP pairing (empty symbols) and the uEPSP burst (filled symbols). (C), The summary data are showing the effect of APV (orange) and subsequent GABA application (green) on the $Ca^{2+}$ transients induced by the bAP/uEPSP pairing (empty symbols) and the uEPSP burst (filled symbols).

The data are presented as mean ± SEM. NS $p > 0.05$, *$p < 0.05$, two-sample *t*-test.
